# Uptake of fucosylated type I human milk oligosaccharide blocks by *Bifidobacterium longum* subsp. *infantis*

**DOI:** 10.1101/2025.02.07.637197

**Authors:** Morten Ejby Hansen, Mikiyasu Sakanaka, Mathias Jensen, Hiroka Sakanaka, Michael Jakob Pichler, Shingo Maeda, Julie Franck Høvring, Aruto Nakajima, Sonja Kunstmann, Tine Sofie Nielsen, Günther Herbert Johannes Peters, Dirk Jan Slotboom, Jens Preben Morth, Takane Katayama, Maher Abou Hachem

## Abstract

Human milk oligosaccharides (HMOs) are uniquely rich in the type 1 building block disaccharide lacto-*N*-biose I (LNB, Galβ1,3GlcNAc), as compared to other mammals. Most HMOs are fucosylated, *e.g.*, α1,2 and α1,4 fucosylations on LNB blocks, resulting in H type 1 (H1) and Lewis a (Le^a^) epitopes, respectively. The dominance of *Bifidobacterium* in breastfed infant guts hinges on efficient uptake of HMOs by specific ATP-binding cassette (ABC) importers. However, molecular insight into uptake of fucosylated LNB blocks is lacking. Here, we analyzed the uptake of LNB and its fucosylated H1 and Le^a^ trisaccharides, as well as the mucin-derived disaccharide galacto-*N*-biose (GNB, Galβ1,3GalNAc) by an ABC importer form the HMO-utilization specialist *Bifidobacterium longum* subsp. *infantis*. Structural analyses and molecular dynamics simulations explained how fucosylated and non-fucosylated LNB forms are recognized with similar affinities by the binding protein of this importer. Strikingly, we showed that two ABC importers confer to the uptake of LNB, while the Le^a^ trisaccharide is efficiently internalized by a single importer in *B. infantis*. Phylogenetic and structural analyses of bifidobacterial ABC-associated binding proteins showed that the Le^a^ clade harbors homologues possessing internal cavities, which allows for the accommodation of branched oligosaccharides. Our work provides unique insight into the evolution and molecular basis of capture and uptake of key HMO and host-derived saccharide blocks, highlighting these compounds as hitherto unexplored candidates for fortification of infant formula.

**Importance:** The assembly of the gut microbiota in early life is critical to the health trajectory of human hosts. Breast feeding selects for a *Bifidobacterium-*rich community, adapted to efficiently utilize HMOs from mother’s milk. Industrial scale production of HMOs for infant formula fortification has mainly considered fucosyllactoses, whereas fucosylated type 1 HMO blocks have hitherto not been explored. Our work sheds light on the uptake facet, central to the utilization of fucosylated HMOs with type 1 LNB building blocks. These type I blocks are efficiently internalized and assimilated by *B. infantis*, which has been recently shown to secrete immune-modulatory aromatic-lactate metabolites that mediate immune-priming of hosts in early life. This study contributes to our understanding of the utilization of HMOs and highlights fucosylated LNB blocks, as hitherto unexplored prebiotic candidates that support the growth of *B. infantis* and other beneficial bacteria in early life.

## INTRODUCTION

The natural assembly of the early life human gut microbiota (GM) is critical for the host’s health trajectory, including the maturation and homeostasis of the immune system (1, 2) as well as the modulation of host metabolism (3). The assembly of the neonate GM is thought to be initiated via orthogonal transfer from the mother during vaginal delivery (4). Thereafter, breastfeeding assumes a prime role in selecting a *Bifidobacterium* dominated gut community (5). Breastfed infants are preferentially colonized by distinct bifidobacteria such as *Bifidobacterium bifidum, Bifidobacterium breve* and the HMO-utilization specialists *Bifidobacterium longum* subsp. *infantis* (6, 7). This specific *Bifidobacterium*-rich signature is associated with protection from immune disorders (8, 9) and enteropathogens (10), which is partially attributed to metabolites, *e.g.*, aromatic-lactates that are secreted by this community, especially *B. infantis* (11). The abundance of this subspecies decreases as solid food dominates the infant food, which highlights its adaptation to human milk oligosaccharides (HMOs) from mother’s milk. The specialization of bifidobacteria in the infant gut is effectuated by numerous genetic loci that mediate the catabolism of HMOs and related host-derived glyco-conjugates (*e.g.*, from mucin) (12–15). HMOs, which are non-digestible by pancreatic enzymes and highly diverse oligosaccharides with a degree of polymerization >3, are the third most abundant component of mother milk (about 12 g L^-1^) (16), thereby constituting the main metabolic resource for the GM of breastfed infants.

The utilization strategy of bifidobacteria involves the import of intact or partially degraded oligosaccharides for subsequent intracellular enzymatic depolymerization and catabolism. The main oligosaccharide importers in bifidobacteria are of the ATP-binding cassette (ABC) type. Specialized extracellular solute binding proteins (SBPs) associated to ABC transporters mediate high-affinity capture of ligands for translocation into the cytoplasm through permease domains (17). Recent studies show that SBPs of bifidobacterial ABC transporters govern saccharide uptake selectivity (18), thereby supporting competitive growth on preferred ligands in mixed cultures (19, 20) and in guts of breastfed infants (21). Despite this pivotal role, bifidobacterial HMO transporters have received little attention. The first molecular study on HMO importer described the structural characterization of GL-BP, the SBP that confers the capture of HMO type I building block lacto-*N*-biose I (LNB, Galβ1,3GlcNAc) in *Bifidobacterium longum* (22). This protein also displays 8.5-fold higher affinity towards the mucin core 1 disaccharide galacto-*N*-biose (GNB, Galβ1,3GalNAc, also called T-antigen) relative to LNB. Another study suggested that at least two SBPs have affinity to LNB/GNB in *B. infantis*: locus tag Blon_2177 (ortholog of GL-BP in *B. longum*) and a divergent protein (Locus tag Blon_0883) that additionally displayed affinity to the blood group H type 1 epitope (H1) and the Lewis a (Le^a^) trisaccharides based on glycan array analysis (15). The contribution of these ABC transporters to HMO and fucosylated host-derived oligosaccharides, however, remains unknown.

Here, we analyzed the capture of GNB as well as non-fucosylated and mono-fucosylated forms of LNB by Blon_0883. We also determined three different complex structures of this SBP bound to GNB, LNB and the H1 trisaccharide, unravelling a different binding site architecture as compared to the previously structurally characterized GL-BP (ortholog of Blon_2177 from *B. infantis*). We demonstrated how this architecture allows the binding of the H1 and Le^a^ epitopes in different conformations by Blon_0883. We also used gene inactivation and uptake assays to evaluate the contribution of the two above-mentioned ABC transporters to the uptake of GNB and LNB blocks. Our study brings novel insight into the function and evolutionary trajectory ABC importers of fucosylated HMO type I building blocks from infant gut bifidobacteria and highlights the relevance of these HMO blocks as candidates for infant formula fortifiers.

## RESULTS

### Blon_0883 binds GNB, LNB and mono-fucosylated LNB trimers with similar affinities

The mature peptide of Blon_0883 was produced and purified to electrophoretic homogeneity. The binding affinity and thermodynamics of the transport protein were analyzed by isothermal titration calorimetry (ITC) and surface plasmon resonance (SPR) (FIG 1, Supplementary FIG S1, Table S1). Among the tested ligands, the highest affinities were measured towards LNB and GNB. The binding of both ligands was driven by favorable enthalpy, similarly to other saccharide-specific SBP (21) (Table 1). The Lewis a (Le^a^) and Blood group H type 1 antigen (H1) trisaccharides were both bound with about 4-fold lower affinity than the preferred LNB ligand. Similar affinities for the Le^a^ and H1 ligands suggest the accommodation of either an α1,2 or α1,4-linked fucosyl unit to either the Gal or the GlcNAc of the LNB backbone, respectively. No binding was observed to lactose or the major fucosyllactose HMOs, underscoring the specificity to the LNB/GNB backbone. Based on these data, Blon_0883 from *B. infantis* is henceforth designated as a GNB and fucosylated LNB-binding protein (*Bi*GFL-BP).

**FIG 1.**
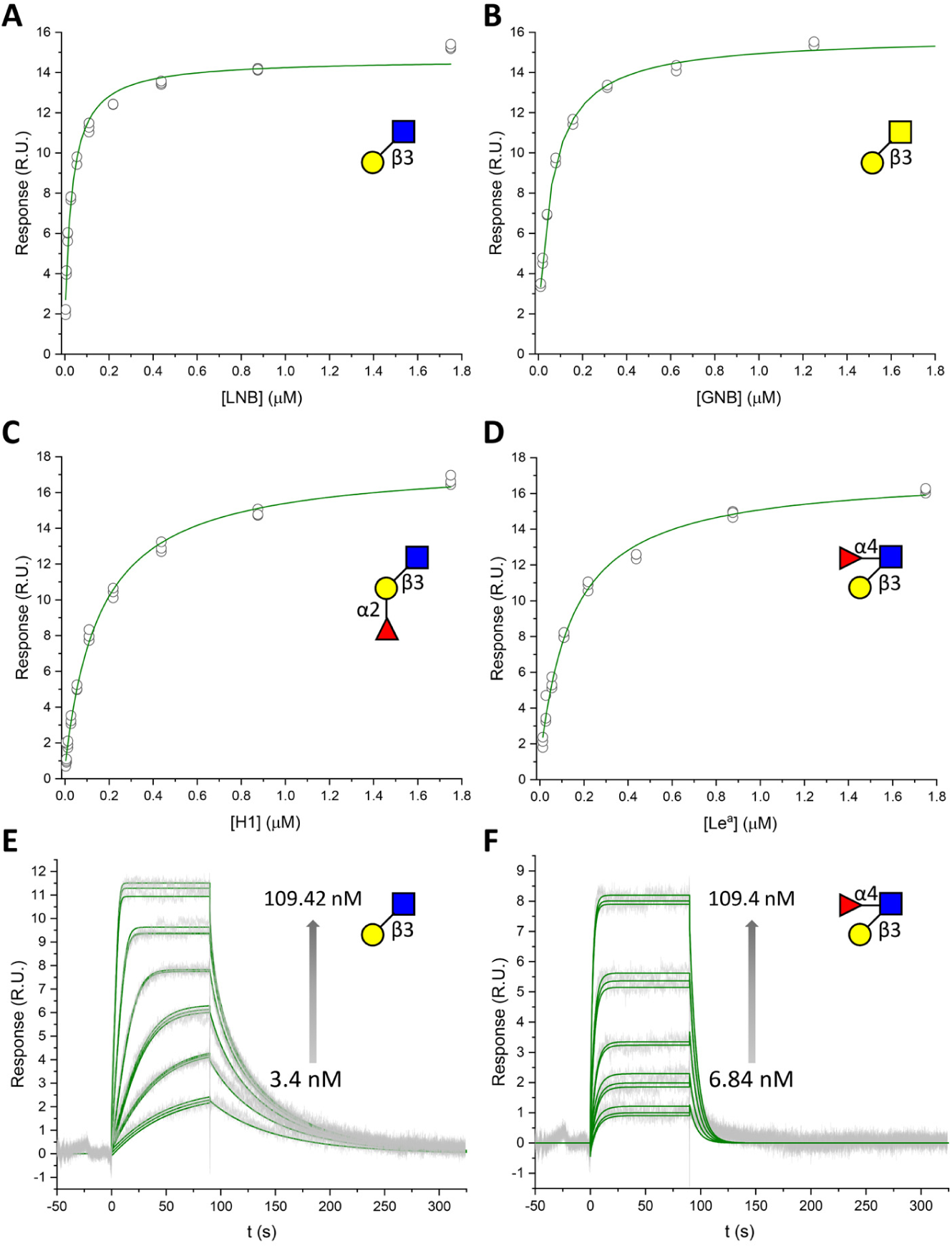
Binding of GNB and LNB as well as its different fucosylated forms to Blon_0883 (*Bi*GFL-BP) as analyzed by surface plasmon resonance at 25°C and pH 6.5. Panels (A)-(D) depict representative binding isotherms based on technical triplicates. The circles are blank and reference-corrected binding levels, and the solid lines are the fits of a one binding site model to the steady state data. Panels E and F depict the blank and reference-corrected sensograms in grey and the green lines are the global fits of a kinetic one site binding model to the binding sensograms.

**Table 1.** Binding parameters of *Bi*GFL-BP determined by isothermal titration calorimetry and surface plasmon resonance analyses, both at 25 ⁰C.

| | $K_a (\times 10^6 \text{ M}^{-1})$ | $K_d (\text{nM})$ | | $\Delta G (\text{kcal mol}^{-1})$ | $\Delta H (\text{kcal mol}^{-1})$ | $-T\Delta S (\text{kcal mol}^{-1})$ | $n$ |
| --- | --- | --- | --- | --- | --- | --- | --- |
| Lacto-*N*-biose | 14.6 ± 3.3 | 68.5 | <sup>A</sup>32.5±3.2 | -9.75 | -13.0 ± 0.3 | 3.25 | 0.92 ± |
| Galacto-*N*-biose | 5.9 ± 0.85 | 169 | <sup>A</sup>62.0±4.0 | -9.18 | -11.0 ± 0.1 | 1.82 | 0.98 ± |
| Le<sup>a</sup> | - | - | <sup>A</sup>146±12 | - | - | - | - |
| H1 | - | - | <sup>A</sup>158±9.0 | - | - | - | - |

<sup>A</sup>The $K_d$ values determined from the steady state SPR analysis.

### Structural rationale for capture of fucosylated LNB forms by *Bi*GFL-BP (Blon_0883)

The crystal structures of *Bi*GFL-BP complexed with LNB and GNB were solved at 2.1 Å and 1.4 Å, respectively (Table S2). *Bi*GFL-BP adopts a canonical SBP fold (Structural cluster B (23)), comprising two domains joined by a tripartite hinge with the ligand binding site located at the domain interface (FIG 2A). Domain 1 (36–158; 327–371) is formed by eight α-helices and five β-strands. Domain 2 (163–322; 376–443) consists of nine α-helices and 7 β-strands, and the hinge region comprises two short loops spanning the center of the two domains (159–162; 323–326) and the loop 373–375. The structures of *Bi*GFL-BP-LNB/GNB complexes reveal well-defined densities for both disaccharides in the binding site (FIG 2B,C). The common galactose (Gal) moiety in both LNB and GNB is recognized by a potential bidentate polar interaction between D210 and the C4-OH and C6-OH groups, establishing the specificity towards a galactosyl unit at this position. Binding of a glucosyl with an equatorial C4-OH is hindered via steric clashes with a glutamine (Q89) that forms a water mediated hydrogen bond to the cyclic oxygen. Other direct polar interactions are provided by the backbone amide and the side chain of T205. Moreover, a water molecule mediates a potential hydrogen bond between the side chain of S164 and the C6-OH. A key aromatic stacking of the Gal unit (both in LNB and GNB) onto Y208 defines position 1, similarly to other binding proteins(19, 24) (FIG 2B). The GlcNAc (in LNB) and the GalNAc (in GNB) are also densely recognized with three potential polar interactions and aromatic stacking onto W290. Thus, the interactions between G47 and E294 are shared between the LNB and GNB complexes, whereas Glu327 recognizes either the C4-OH in the GalNAc unit of GNB or either the C4-OH or/and the C6-OH of the GlcNAc unit in LNB via hydrogen bonding (FIG 2B,C). Interestingly, a partially solvent accessible cavity large enough to accommodate a monosaccharide was observed facing the C2-OH of the Gal moiety (FIG S2), whereas the region around the C4-OH of the GlcNAc in LNB is less spacious. To bring insight into binding of fucosylated LNB forms, we attempted to co-crystallize *Bi*GFL-BP with a commercial preparation of Le^b^, which resembles a superimposition of H1 and the Le^a^ trisaccharides to assess whether the protein can accommodate both fucosylations simultaneously. Surprisingly, the structure of the co-crystallized *Bi*GFL-BP, revealed electron density for the α-1,2-linked fucose (Fuc) to the Gal unit of LNB, *i.e.* we got the structure of the protein in complex with the H1 trisaccharide. This indicated that binding to the Le^b^ tetrasaccharide was too weak or unfeasible, relative to the H1 trisaccharide, which maybe have been a contaminant of the commercial preparation. The structure of the LNB backbone in the H1-complex overlays with LNB in the LNB-complex and the fucosyl unit was bound in the above mentioned partially solvent accessible cavity above the Gal unit of LNB (FIG 2B,D and FIG S2). A single hydrogen bond from the side chain of T101 to the C3-OH of the fucosyl unit was observed. Notably, the methyl group of the fucosyl unit is accommodated by an apolar pocket formed mainly by the F44 and S100 and the acetyl group of the GlcNAc in LNB. The electron density of the fucosyl is less well defined than that of the LNB backbone, in line with a single polar interaction, allowing for partial flexibility of this sidechain.

**FIG 2.**
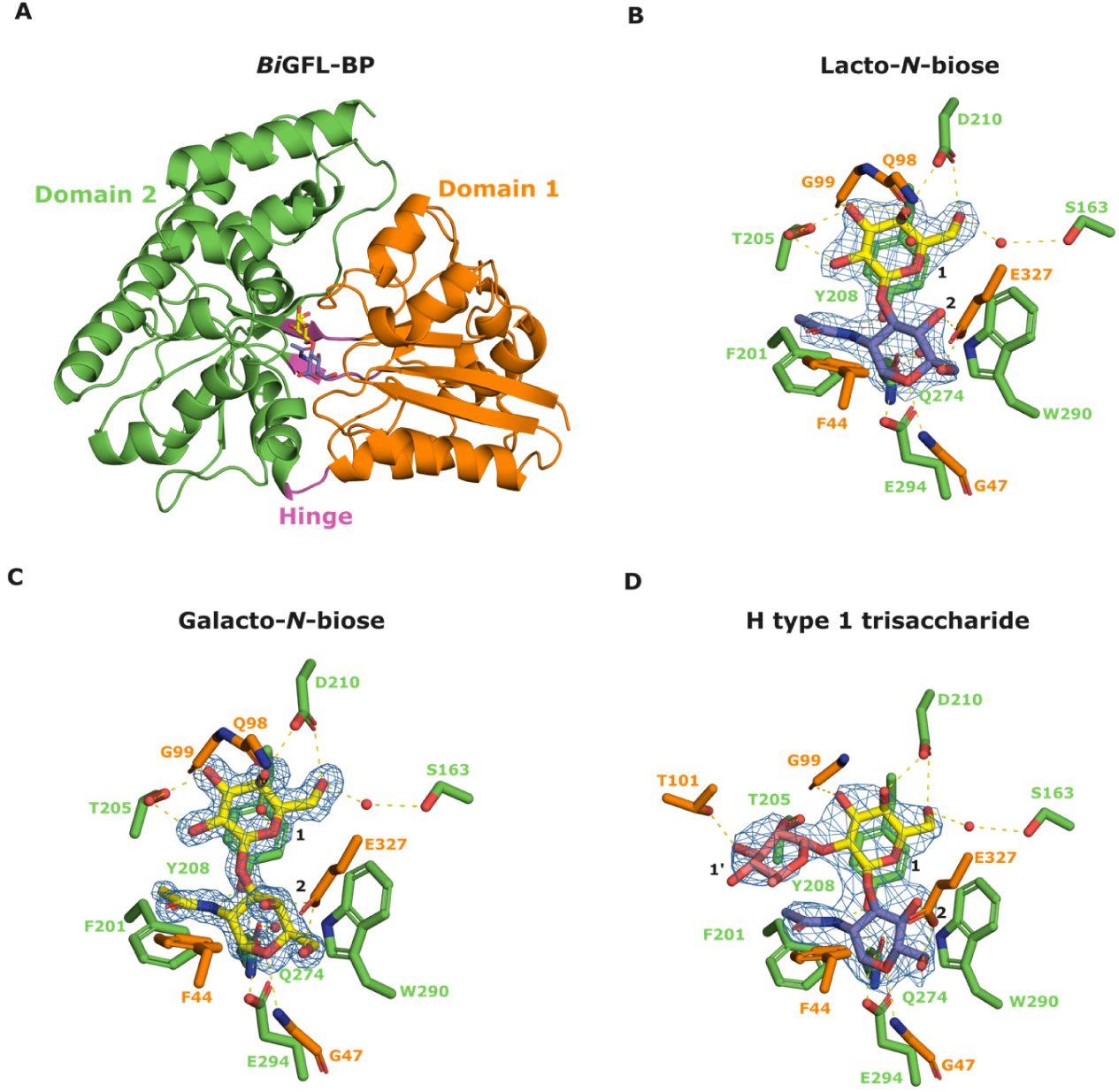
The overall structure of *Bi*GFL-BP and the binding site interactions with ligands. (A) Overall structure of *Bi*GFL-BP in complex with LNB. (B), (C) and (D) show the binding details of *Bi*GFL-BP to LNB, GNB and the H1 trisaccharide, respectively. The binding residues are colored according to domains presenting them. Water molecules are denoted with red spheres and potential polar interactions are denoted as dashed yellow lines. The electron density map (2F_O_-F_C_), contoured at 2σ, is shown as a blue mesh around the ligands.

### Molecular dynamics simulations

To explore the mode of binding of the Le^a^ trisaccharide, we resorted to molecular dynamics (MD) simulations. We started by performing the MD simulations based on the H1-complex to study the dynamics and key interactions that confer binding with *Bi*GFL-BP. The protein exhibited a marked degree of fluctuation, as also reflected in root mean square deviations (RMSD), with plateaus of 2-4 Å (FIG S3). In addition, the root mean square fluctuations (RMSFs) were qualitatively comparable to the B-factor determined from the crystal structure (FIG S4). Despite these fluctuations, prominent interactions, identified over the course of the simulations, were in accordance with the crystal structure complex with the same ligand. A polar interaction between D210 and the hydroxyl group at C4 (C4-OH) in the Gal unit is observed as well as aromatic staking of Gal onto Y208. In addition, W290 forms stable polar interactions with and aromatic stacking onto the GlcNAc unit. Overall, the most persistent interactions occur between the protein and the LNB backbone of the H1 trisaccharide ligand. Accordingly, the largest contribution to the binding affinity is provided by these two sugar moieties, consistent with the binding experiments and the ligand-bound crystal structure. With this validation, we wanted to gain insight into the binding of the Le^a^ trisaccharide. Since the mode of binding of this ligand was unknown, we performed MD simulations for two docked conformers of the Le^a^ trisaccharide. The binding energies of the ligands were estimated using the molecular mechanics generalized Born surface area approach in CPPTRAJ (see material and methods). The binding energies of both binding modes of the Le^a^ ligand are comparable to the binding affinity of the H1 trisaccharide (Table S3), suggesting that the binding of the Le^a^ ligand may be feasible in either orientation. A comparison of the simulations of the two binding modes and the simulations of the H1-complex reveals that forementioned interactions with Y208, D210, and W290 are consistently present and stable across all simulations (FIG 3). This indicated that these interactions play a crucial role in the binding of fucosylated LNB ligands, similarly to the non-fucosylated disaccharide. Notably, an additional interaction between the Gal unit in the Le^a^ ligand and S163 was observed in the simulations, whereas a solvent mediated hydrogen bond was observed in the crystal structure of the H1 complex. Comparison of the two putative binding modes of the Le^a^ trisaccharides showed that the overlay of the LNB backbone was worse for confirmer 1, where the fucosyl is accommodated in the less spacious internal cavity of the protein (FIG 3A, FIG S5). By contrast, this overlay was better for conformer 2, where the fucosyl unit was bound in the same fucosyl-binding cavity in the H1 complex structure (FIG 3B, FIG S5). Altogether, these data were consistent with the binding of Le^a^ by the protein. Differences in conformations and positioning of the LNB backbone are likely to hamper the double fucosylated Le^b^ backbone, as hinted by the soaking experiment. This was confirmed experimentally using an uptake assay that showed that the Le^b^ tetrasaccharide was not internalized after 6 h of incubation with *B. infantis* cells (FIG S6).

**FIG 3.**
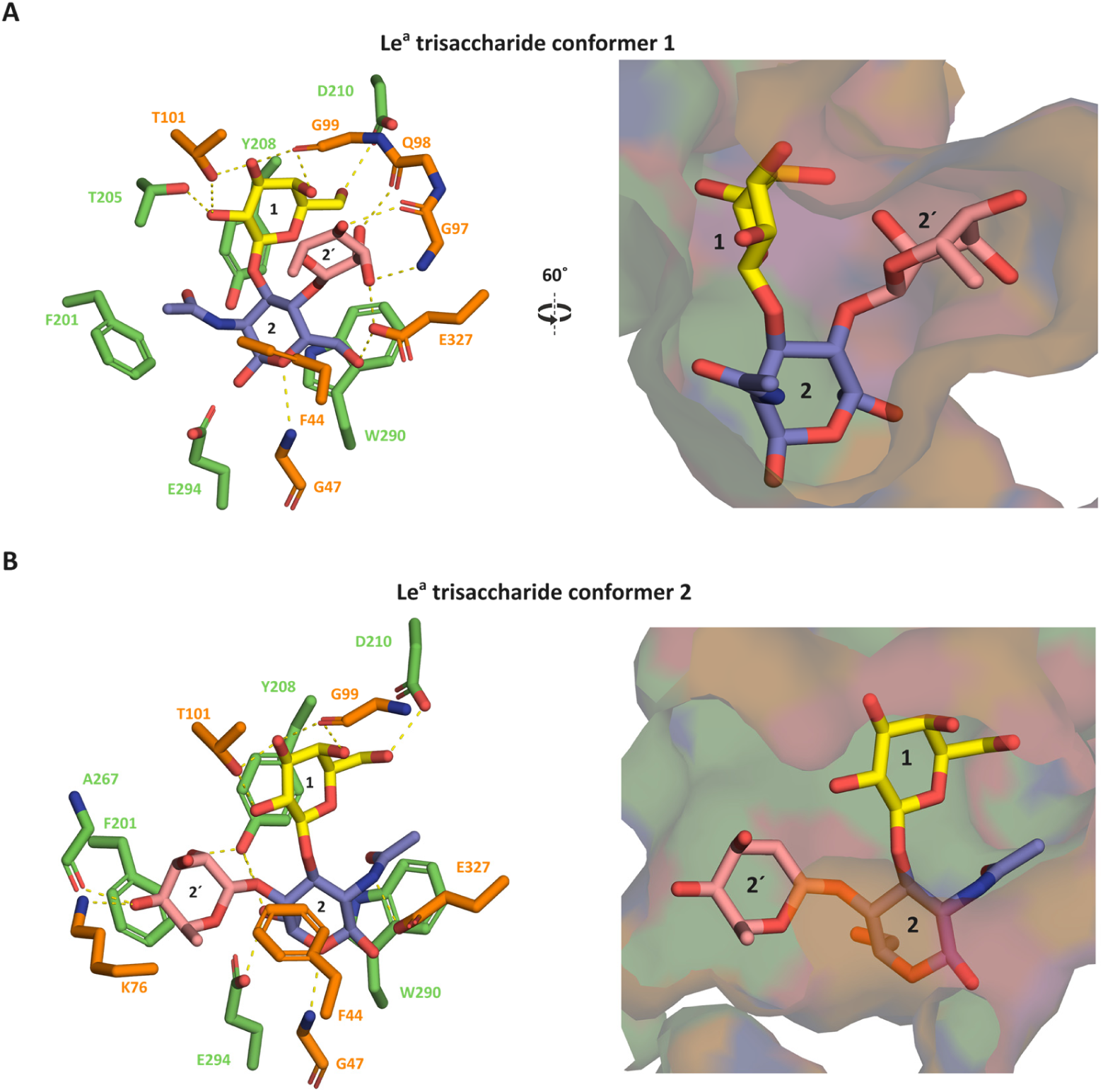
Molecular dynamic simulations of the Le^a^ trisaccharide binding to *Bi*GFL-BP. (A) and (B) depict two modes of accommodation of the Le^a^ trisaccharide in the binding site of *Bi*GFL-BP. Amino acid residues participating in the binding of the ligand are colored according to the domain they originate from and the galactose, *N*-acteylglucosamine and fucose units of the ligand are colored in yellow, blue and salmon, respectively. The yellow dots in the left panels denote potential hydrogen bonds. The left panels are semi-transparent surface representations of the binding cavities that accommodate the fucosyl unit of the ligand. The conformer 1 shown in (A), the fucosyl unit is positioned into a tight cavity, whereas flipping the *N*-acteylglucosamine unit of LNB positions the fucosyl in the more spacious cavity, similarly to the H1 trisaccharide shown in FIG 2.

### *Bifidobacterial* HMO-specific transport binding proteins have evolved from different ancestors

To compare *Bi*GFL-BP to structurally described homologs, we performed A DALI structural similarity search (25) against the Protein Data Bank. This search identified the galactose binding SBP from *Actinoplanes* sp. (PDB ID: 3OO6-A, Z-score=33.3), the verbascose SBP from *Streptococcus pneumoniae* (PDB ID: 6pre-C, Z-score=32.6) and the chitooligosaccharide SBP from *Paenibacillus* sp. FPU-7 (PDB ID: 7ehp-B, 32.6) as the three top structurally related hits with shared sequence identities of 15, 20 and 21%, respectively, to *Bi*GFL-BP.

Despite the common structural fold and the shared LNB/GNB-binding function, *Bi*GFL-BP and the previously described GL-BP from *B. longum* (PDB code: 2Z8F) (22) share low structural similarity (Z-score=25.5; overlay RMSD =3.4 Å for 49 aligned C_α_ atoms). The divergence of these proteins is underscored by the entirely different binding architecture, with the LNB ligand being bound in an almost orthogonal orientation relative to each other (FIG 4A). Strikingly, the structurally closest bifidobacterial ortholog to *Bi*GFL-BP is the arabino-xylooligosaccharide binding protein from *Bifidobacterium animalis* subsp. *lactis* BL-04 (*Bi*AXOS, PDB ID: 4c1u, Z-score=32.2, RMSD =3.3 Å for 363 aligned C_α_ atoms, 17% sequence identity) (24). Interestingly, the binding directionality of the saccharide backbone and the position of the cavity accommodating the fucose/arabinose side chains in these two SBP are similar (FIG 4B).

**FIG 4.**
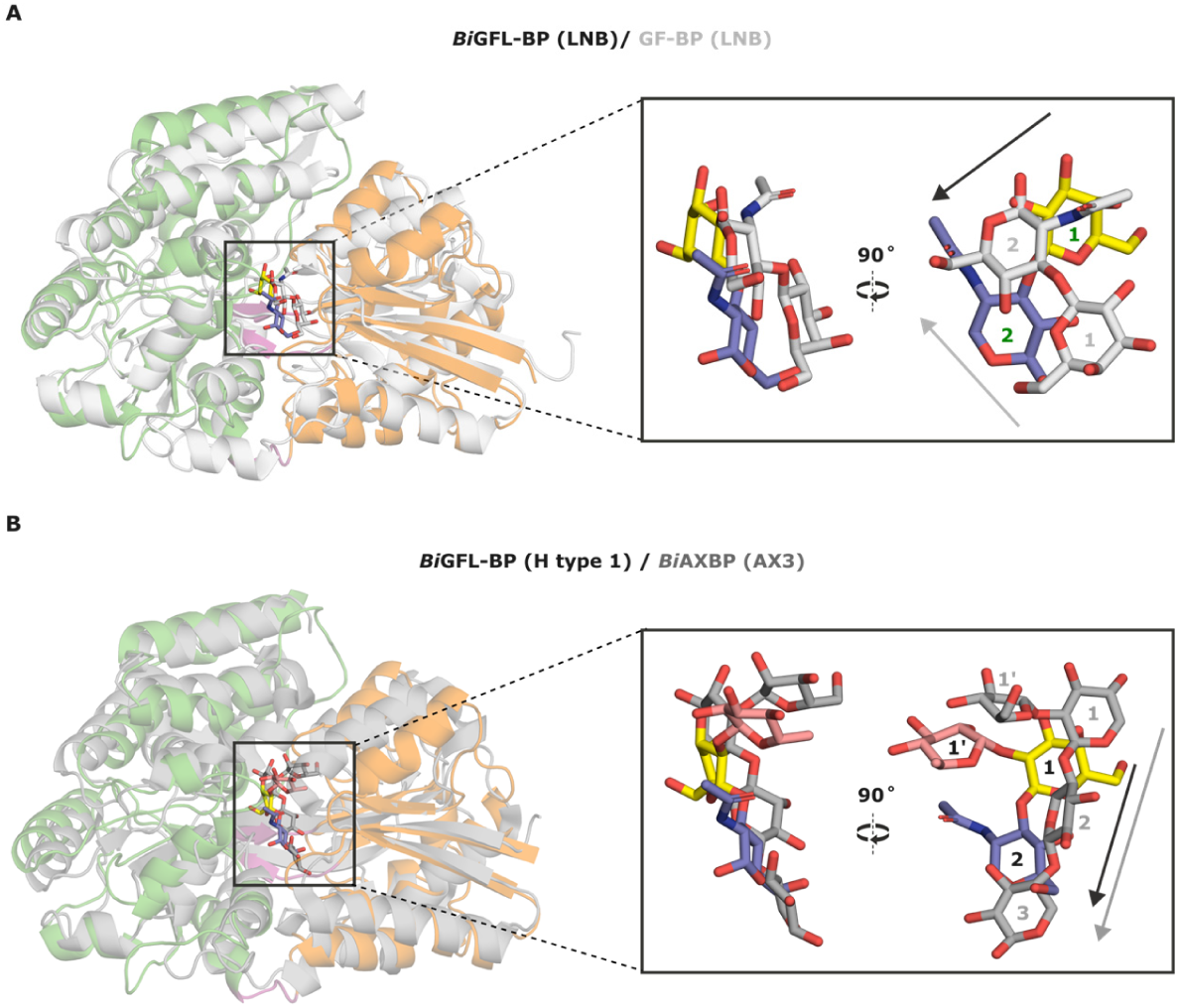
Comparison of binding site architectures between *Bi*GFL-BP and structural homologs. (A) Structural overlay between *Bi*GFL-BP described in this study and the previously characterized GNB/LNB binding protein from *Bifidobacterium longum* in complex with LNB (GL-BP, PDB ID: 2Z8E). (B) A structural overlay of *Bi*GFL-BP with the arabino-xylooligosaccharide binding protein from *B. animalis* subsp. *lactis* in complex with arabinoxylotriose (PDB: 3ZKL). *Bi*GFL-BP was colored according to domain. The superimpositions were performed in PyMOL v. 2.5.5 (RMSD 3.85 Å for 1701 aligned atoms for A and 3.02 Å for 1834 aligned atoms for B, respectively).

To explore the evolutionary relationship between the HMO-specific SBPs, we performed a phylogenetic analysis of a total of 379 *Bifidobacterium* SBP sequences, based on all biochemically and structurally characterized solute binding proteins. The phylogenetic tree revealed high divergence of the sequences (FIG 5). Most characterized HMO-specific SBPs, *e.g.*, fucosyllactose, tetrasaccharide lacto-*N*-tetraose (LNT, Galβ1,3GlcNAcβ1,3Galβ1,4Glc), and GNB/LNB binders, populate two large clusters. Remarkably, *Bi*GFL-BP segregates from the above HMO binding proteins and clusters in a branch adjacent to the arabino-xylooligosaccharide binding protein from *B. animalis* subsp. *lactis*, underscoring the architectural similarity of these SBPs. Another *B. infantis* SBP (Blon_2343), displaying binding to HMOs containing a type II building block *N*-Acetyllactosamine (LacNAc, Galβ1,4GlcNAc) on glycan arrays (15) is also within the same cluster as *Bi*GFL-BP. This is consistent with the structural similarity of *Bi*GFL-BP to the chitooligosaccharide binding protein from *Paenibacillus* (*vide supra*), which may stem from the recognition of a β-glycosidic bonds and bulky *N*-acetylamine groups in these ligands by these three SBPs. In summary, these findings suggest two evolutionary origins of *Bi*GFL-BP and GL-BP. The structural similarity between *Bi*GFL-BP and *Bi*AXOS indicates that this architecture has evolved from a common ancestor to accommodate backbones with variable branching (*e.g.*, fucose or arabinose), which could fit in a spacious cavity positioned above the main chain binding site.

**FIG 5.**
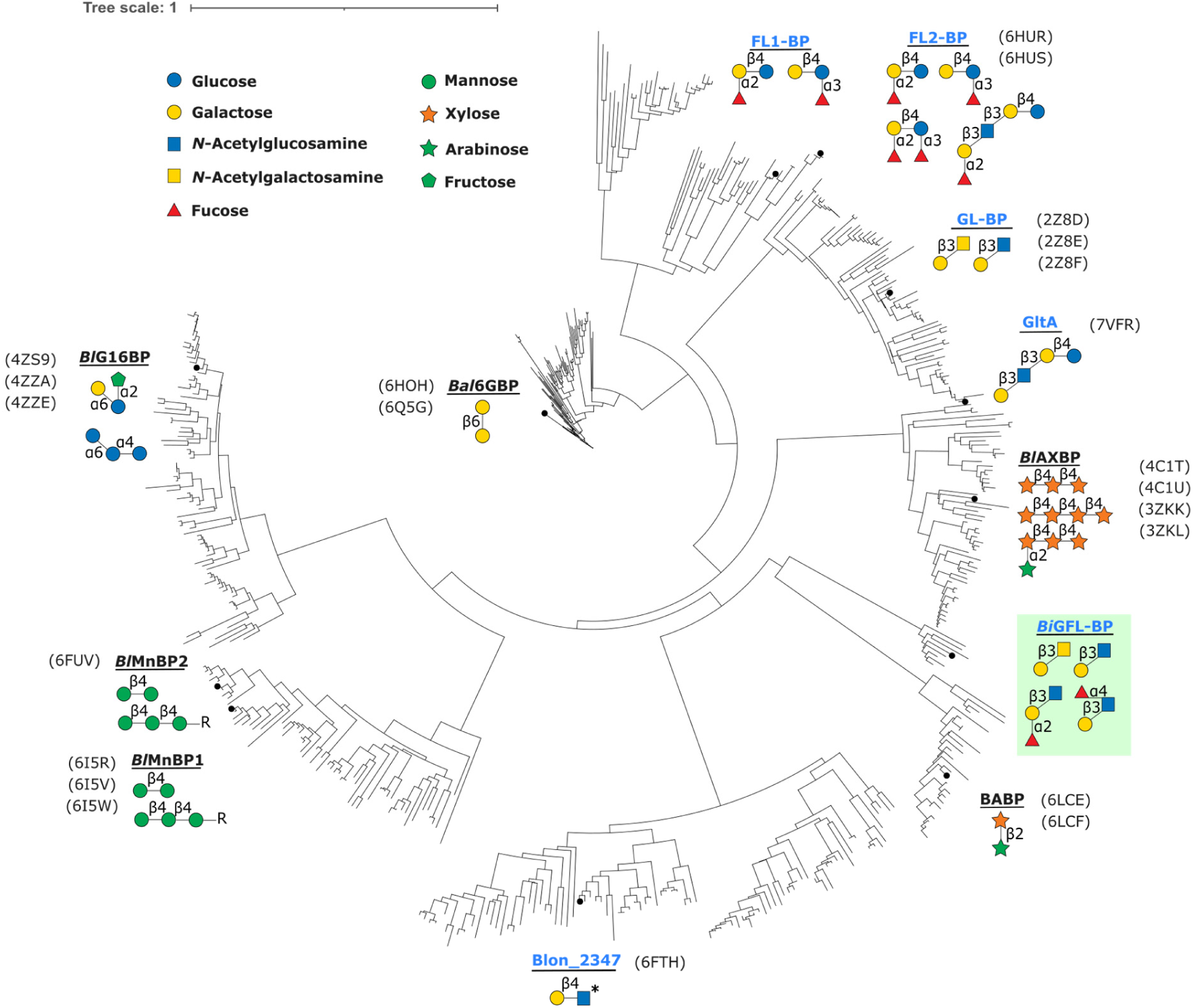
Phylogenetic analysis of solute binding proteins from *Bifidobacterium*. The tree was computed based on 379 ABC-associated binding proteins from *Bifidobacterium* sharing an amino acid sequence identity >30%. Structurally and/or biochemically characterized are depicted on the tree as follows: *Bal*GBP (18), β(1,6)-galactoside binding protein from *B. animalis* subsp. *lactis* Bl-04; FL1-BP and FL2-BP, fucosyllactose and other fucosylated HMO binding proteins from *B. infantis* (21); GltA (Blon_2177); Structure of a *B. infantis* SBP in complex with the HMO lacto-*N*-tetraose (LNT, PBD ID: 7VFQ); GL-BP (66), Lacto-*N*-biose binding protein from *B. longum* JCM1217; *Bl*AXOS (24), arabino-xylooligosaccharide binding proteins from *B. animalis* subsp. *lactis* Bl-04; *Bi*GFL-BP (Blon_0883, green highlight), the solute binding protein described in the present study; BABP (67), β-_L_-arabinobiose-binding protein from *B. longum*; Blon_2347 (15), *B. infantis* SBP found to be bind to lactosamine motifs in a glycan array analysis; BlG16BP (19), α-(1,6)-linked galacto/glucooligosaccharides from *B. animalis* subsp. *lactis* Bl-04. *Bl*MnBP1/2 (20), β-mannooligosaccharide binding proteins from *B. animalis* subsp. *lactis* Bl-04. The HMO specific binding proteins are highlighted in blue.

### The ABC transporter associated to *Bi*GFL-BP is the main uptake route for the Le^a^ trisaccharide

To evaluate the contribution of the ABC transporter associated to *Bi*GFL-BP (Blon_0883), we disrupted the encoding gene in the wildtype strain by a suicide plasmid-mediated single-crossover event. The *Bi*GFL-BP gene disruption was also performed in a mutant strain where the entire ABC transporter genes associated to the GL-BP ortholog (Blon_2175–2177) was deleted (Sakanaka et al., in revision and materials and methods) (FIG S7, Table S5). The resulting GL-BP, *Bi*GFL-BP, and double knockout mutants were used for the growth experiments using different sugars as sole carbohydrate sources. As expected, the wildtype and mutant strains grew indistinguishably on galactose, consistent with galactose not being a substrate for either ABC importer (FIG 6A). Both the wildtype and single mutants grew equally well on LNB. Further, this disaccharide sustained the growth of the double mutant, albeit less efficiently than the wildtype strain. Thus, the growth of the double mutants on LNB was reduced to about a third of the value of control, although either single mutant had a modest effect on growth. These findings establish that the two importers analyzed are the main uptake routes for LNB in *B. infantis*. These conclusions are also supported by the high residual LNB concentration in the supernatant of the culture of the double mutant strain (74% of the total) as opposed to the wildtype and the single mutants (FIG 6B). This apparent multiplicity underscores the importance of this building block for fitness in the breast-fed infant gut. By contrast, the double mutant grew strongly on GNB, revealing that efficient uptake of GNB can be mediated by additional transporters.

**FIG 6.**
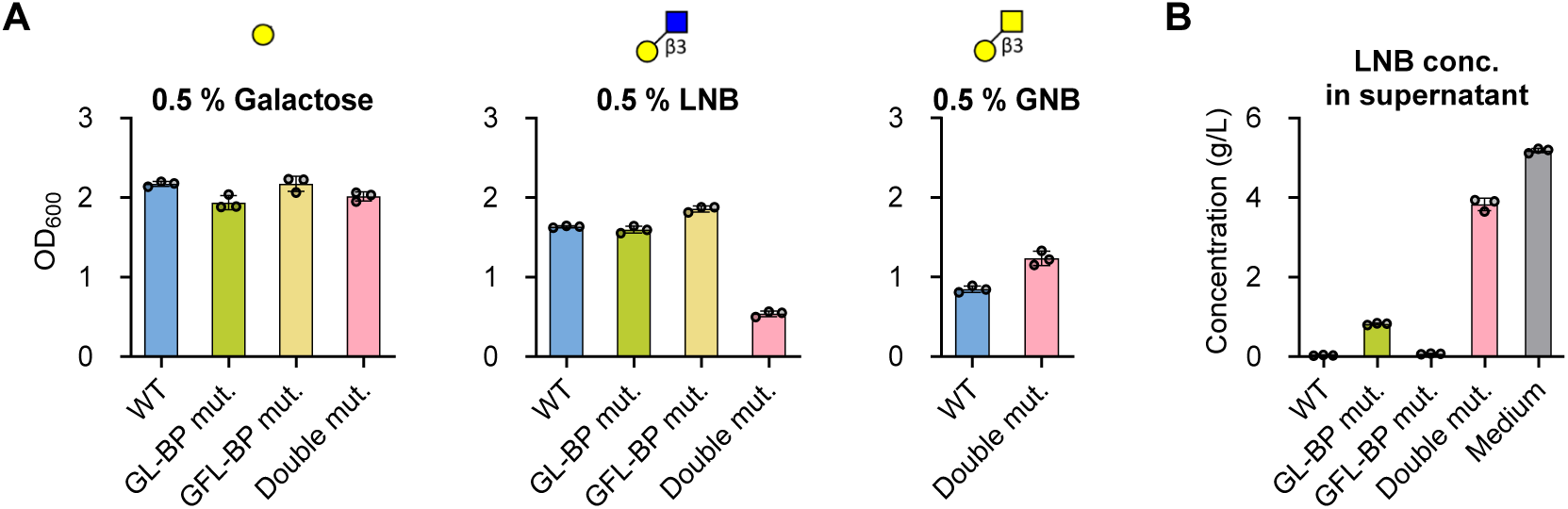
Uptake of LNB and GNB by *B. infantis* gene knockouts of GL-BP and *Bi*GFL-BP. A) Growth of *B. infantis* wild-type (WT) and GL-BP– and *Bi*GFL-BP-single and double gene knockout strains in the media supplemented with 0.5 % (w/v) Gal, LNB, or GNB. *OD*_600_ was measured post 12 h (Gal and LNB) and 18 h (GNB) of cultivation. B) LNB concentrations in the spent media at 12 h of incubation. Data are represented as dot plots with mean ± SD of biological triplicates.

The uptake of Le^a^ and H1 trisaccharides by *B. infantis* was also examined by monitoring the residual ligand concentration in culture supernatants with thin layer chromatography (TLC) analysis. Both Le^a^ and the H1 antigen trioses appear to sustain the growth of *B. infantis* based on the depletion of these ligands from the culture supernatant of the wildtype strain after 5 and 9 hours, respectively (FIG 7). Strikingly, the single *Bi*GFL-BP mutant strain impaired the uptake of Le^a^ (Fig 7A), which indicates that the ABC importer associated to *Bi*GFL-BP is the only efficient uptake route for this substrate. By contrast, the uptake of the H1 triose was largely impaired in the double mutant strain, and to a less extent in the single mutant strain (FIG 7B), indicating that both transporters contribute to the uptake of this trisaccharide. These data clearly suggest that the two ABC systems collectively make an important contribution to the uptake of LNB and H1 trisaccharide, but that the *Bi*GFL-BP associated ABC transporter is critical for the uptake of the Le^a^ trisaccharide.

**FIG 7.**
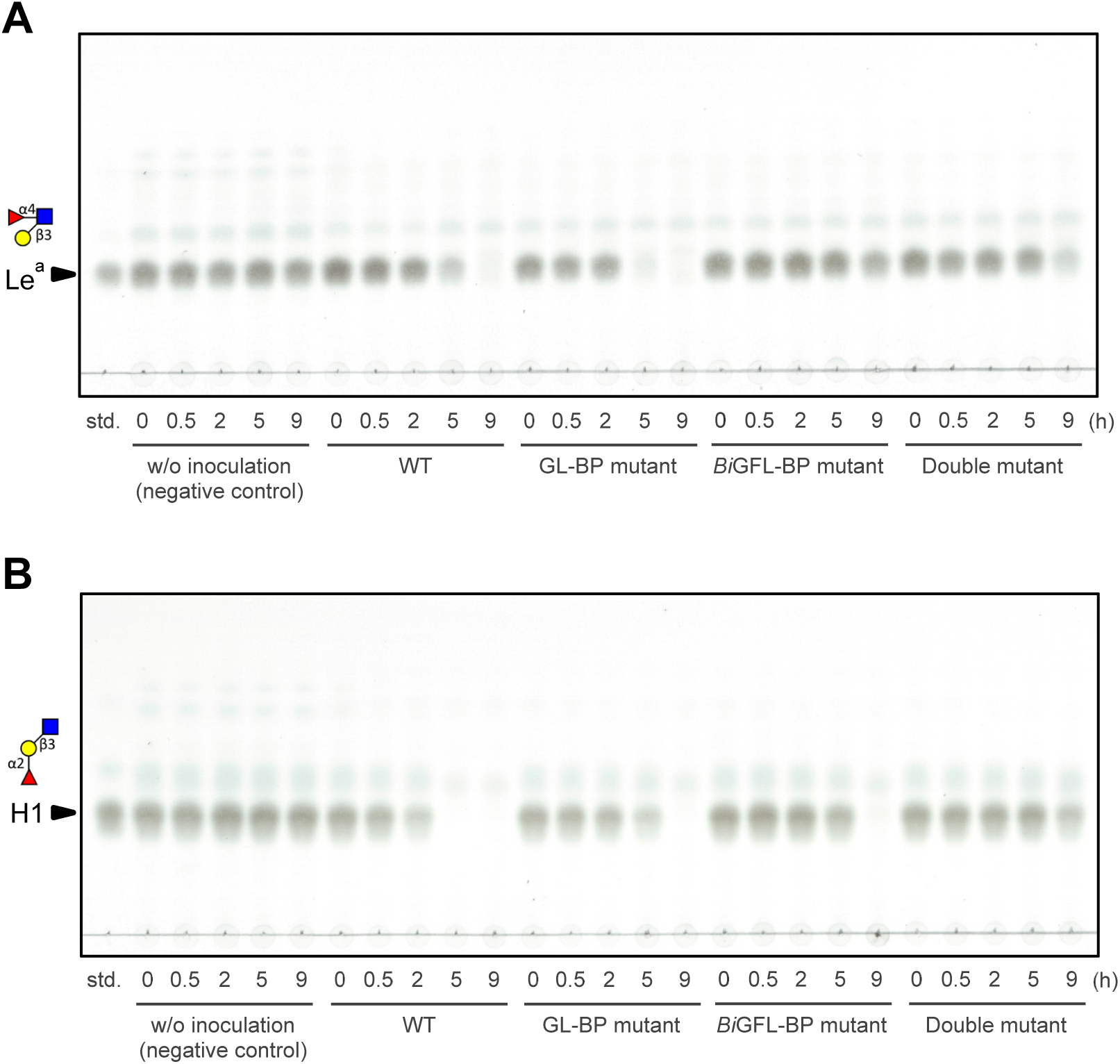
Contribution of *Bi*GFL-BP to the uptake of Le^a^ and H1 trisaccharides. (A, B) WT and mutant cells of *B. infantis* were incubated in the presence of 5 mM Lewis a triose (Le^a^) (A) and blood group antigen type-1 triose (H1) (B). Samples were taken at the indicated time points. Culture supernatants were analyzed by thin-layer chromatography as described in the methods section. Data are representative of biological duplicates.

## DISCUSSION

Compelling evidence supports the crucial rule of HMOs in selecting for a *Bifidobacterium* dominated community in early life, which has life-long impacts on host health. Of the >200 identified HMO structures about 10 exhibit high abundance (16, 26). These include fucosylated and non-fucosylated forms of the tetrasaccharide LNT, comprising an LNB unit joint to lactose via a β(1,3)-linkage. Both *B. infantis* and other *B. longum* strains (27) are key HMO utilizing species in the neonate infant gut. *B. infantis* possesses two ABC importers of the major fucosyllactose HMOs, which were correlated to the abundance of *Bifidobacterium* spp. *in vivo* (21). The less prevalent *Bifidobacterium catenulatum* subsp. *kashiwanohense* and *Bifidobacterium pseudocatenulatum*, possess closely related (> 60 % amino acid sequence identity) homologs to the *B. infantis* fucosyllatose SBPs (28). However, the *B. kashiwanohense* SBP, also encoded by distinct *B. longum* strains, was shown to mediate the uptake of fucosylated LNT HMOs. Different enzymatic routes for the degradation of LNT have been identified. An extracellular GH20 lacto-*N*-biosidase from *B. bifidum* (29) was the first reported enzyme, which catalyzes the release of LNB from HMOs *e.g.*, LNT (FIG 8). An unrelated extracellular lacto-*N*-biosidase from *B. longum* was then reported as the founding member the CAZy family GH136 (27) (FIG 8). The convergent evolution of *B. longum* and *B. bifidum* lacto-*N*-biosidases from unrelated structural scaffolds showcases the importance of harnessing the LNB building block by early life bifidobacteria.

**FIG 8.**
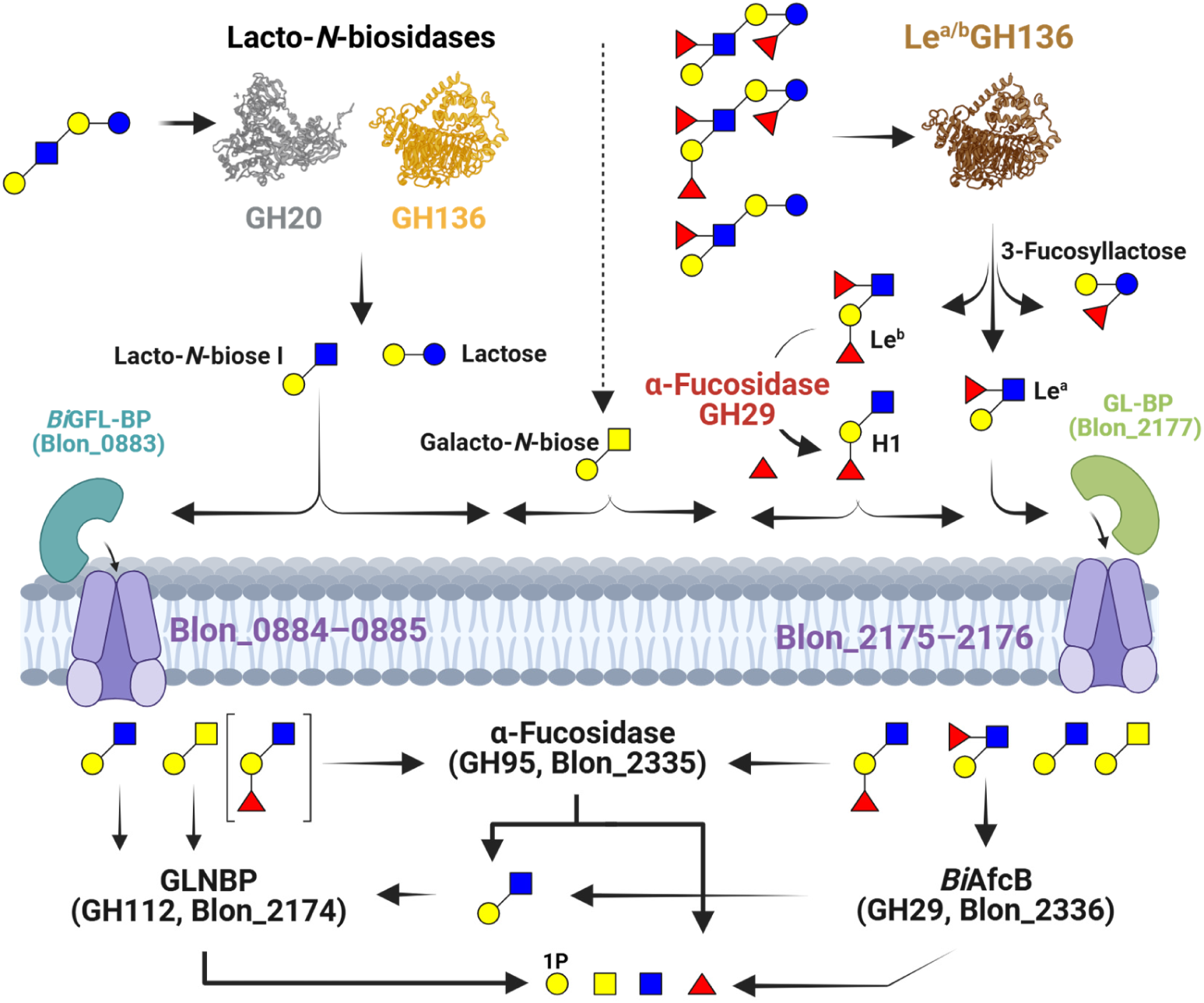
Schematic model for the catabolism of type I HMOs. The figure shows plausible routes of degradation by biochemically characterized enzymes. The specificities and the GH families of extracellular enzymes that can degrade both LNB and its fucosylated forms are indicated. These activities collectively, release LNB, Le^a^ and Le^b^ in addition to lactose and its fucosylated forms. Cleavage of the α1,4-fucosyl unit from the Le^b^ tetrasaccharide liberates the H1 trisaccharide. The specificities of the two ABC transporters, which were shown in the present work to confer the uptake of LNB and its fucosylated forms, besides GNB, are indicated. The square brackets around the H1 trisaccharide, indicate a less preferred uptake route. The intracellular biochemically proven degradation reaction of the internalized substrates by *B. infantis* enzymes is shown. Intracellular enzymes are represented by both their locus tag number in *B. infantis*, enzyme name and GH family. Symbol Nomenclature for Glycans (SNFG) (https://www.ncbi.nlm.nih.gov/glycans/snfg.html), similarly to FIG 1 and FIG 5-7. This figure is made using Biorender.com.

The degradation of fucosylated forms of LNT remained a conundrum until a previously unknown specificity was reported by an extracellular GH136 enzyme from *Roseburia inulinivorans*, a butyrate producing *Clostridium* XIVa member and early colonizer of the human gut (30). This enzyme requires an α1,4-fucosylation at the GlcNAc unit of LNB and tolerated additional fucosylations on the LNT backbone to release the fucosylated LNB forms, *viz.* Le^a/b^ (FIG 8). The released Le^b^ by GH136 (or similar activities) can be further cleaved by the extracellular α-fucosidase (BbAfcB, GH29) from *B. bifidum* (31) or other enzymes to generate the Le^a^ trisaccharide substrate for the ABC system associated to *Bi*GFL-BP that has at least a 10-fold higher affinity than the corresponding binding protein from *R. inulinivorans*, suggesting competitiveness in the capture of this ligand. The evolution of the *Bi*GFL-BP is likely an adaptation to the availability of the H1 and Le^a^ trisaccharides in the infant gut during breast feeding and potentially during the weaning period, when Clostridia such as *R. inulinivorans* likely colonize the infant gut (32). Interestingly, the present study shows that the two ABC transporters associated to *Bi*GFL-BP and the GL-BP from *B. longum* (22) define distinct HMO uptake profiles of LNB HMO blocks. The structural divergence of these two SBPs is another example of convergent evolution, whereby both transporters internalize LNB, but only *Bi*GFL-BP confers efficient capture of Le^a^ and to largely the H1 trisaccharides. Indeed, the gene inactivation data suggested that the GL-BP-associated ABC transporter (Blon_2175-2177) was less efficient in the uptake of the H1 trisaccharide, indicating a lower affinity to this ligand, which underscores the *Bi*GFL-BP-associated ABC transporter as the primary uptake route to the fucosylated LNB blocks (FIG 8). The internalization of fucosylated LNB versions to the cytoplasm is followed by defucosylation and phosphorolysis in *B. infantis*, which possesses GNB/LNB phosphorylase (GLNBP) that has been shown to cleave both these disaccharides to Gal-1-phosphate and GalNAc or GlcNAc in both *B. longum* (12, 27) and *R. inulinivorans* (30) (FIG 8). These diverse uptake and degradation routes of (fucosylated)-LNB blocks are consistent with the adaptation of early life *Bifidobacterium* and other beneficial microbiota members to target these abundant HMO blocks.

In summary, this study highlights key facets in the utilization of HMOs with fucosylated-LNB building blocks, *via* connecting the enzymatic extracellular release by a newly discovered activity assigned into a branch of GH136 enzymes, to uptake through an atypical ABC transporter evolved mainly in *B. infantis* that has high fitness to colonize the breastfed neonate gut. Remarkably, the metabolic capabilities of *B. infantis* and other bifidobacteria to harness abundant HMOs with a fucosylated LNB backbone have received little attention, especially with regard to being excellent substrates for both bifidobacteria and other beneficial early life microbiota members including butyrate producers, proposed to be have an important role in immune homeostasis (33). Current fortification strategies of infant formula have focused on fucosyllactoses as the main HMO supplement. It is important to consider LNB and its fucosylated forms for the fortification of infant formula, and this merits further investigation of the *in vivo* impact of this group of HMO blocks.

## MATERIALS AND METHODS

### Chemicals

The carbohydrates Galactose (Gal), Lactose (Lac), 2’-fucosyl lactose (2FL), 3-fucosyl lactose (3FL), 3’siayl-lactose (3SL), di-fucosyl lactose (DFL) and lacto-*N*-tetraose (LNT) were from were from Sigma-Aldrich (MO, USA). LNB and GNB, both > 95% purity were gifts from Prof. Motomitsu Kitaoka(34, 35) (Niigata University, Japan). *N*-Acetyl-lactosamine (LacNAc), Lewis a triose (Le^a^), Lewis b tetraose (Le^b^) and Blood group antigen H1 triose were purchased from Elicityl (Crolles, France).

### Expression and purification of recombinant Blon_0883

The gene fragment that encodes the mature peptide of Blon_0883 (aa amino acids 24–460) was PCR-amplified and inserted into NcoI and EcoRI sites of pETM11 (a kind gift from Günter Stier, EMBL, Center for Biochemistry, Heidelberg, Germany)(36). The primers used are listed in Supplementary Table S5. The recombinant protein was produced in *E. coli* BL21 (DE3) as an N-terminal fusion with a TEV (Tobacco Etch Virus nuclear-inclusion-a endopeptidase)-cleavable His tag separated by a 3 amino acid insertion (GAM). Protein production was performed by growing the cells in LB medium containing kanamycin at 37 °C to *OD*_600_ = 0.5, thereafter the temperature was reduced to 21 °C and the expression was induced by addition of IPTG to 0.1 mM and growth was continued for 16 h. Cells were harvested by centrifugation, resuspended in binding buffer (10 mM HEPES, 500 mM NaCl, 10 mM imidazole, 10 % glycerol (v/v), 0.5 mM dithiothreitol, pH = 7.4) and lysed by a single passage through a high pressure homogenizer. After centrifugation, clarified lysates were applied onto a 5-mL HisTrap HP column (GE Healthcare, Uppsala, Sweden) and purified as recommended by the manufacturer. Eluted pure fractions were pooled, concentrated as described above, applied to a HiLoad Superdex G75 26/60 gel filtration column (GE Healthcare) and eluted with 10 mM MES buffer (pH 6.5) at 1 mL min^-1^. The His-tag was cleaved using TEV protease as previously described (20). Cleaved proteins were recovered after passing through a HisTrap HP column (1 mL) pre-equilibrated with the binding buffer, concentrated as described above, and stored at 4 °C until further use. Purity was assessed by SDS-polyacrylamide gel electrophoresis. The protein concentration was determined by measuring absorbance (*A*_280_) and using the theoretically calculated (https://web.expasy.org/protparam/) absorption coefficient of 72770 M^-1^cm^-1^ for Blon_0883 calculated based on the amino acid sequence.

### Binding analyses

*ITC analysis−*Binding of Blon_0883 to carbohydrate ligands was analysed by MicroCal iTC_200_ (Malvern, Worcestershire, UK). Proteins (23–25 μM) were dialyzed against 2×500 volumes of 10 mM sodium phosphate buffer, pH 6.5 and thereafter titrated with different ligands (0.25 mM) using 20 injections (0.5 μL for the first injection, 2.0 μL for the following injections separated by 180 s) at 25°C. The ITC thermograms were corrected for the heat of dilution (measured by injecting the ligands into the buffer) and analyzed using the MicroCal Origin 7.0. A binding model for one set of equivalent sites was fitted to the data to determine the equilibrium association constant (*K*_a_), molar binding enthalpy (Δ*H*), and binding stoichiometry (*n*). The experiments were performed in independent duplicates.

*SPR analysis–* Affinity of Blon_0883 to oligosaccharides was also determined using a Biacore T100 (GE Healthcare). The protein was diluted into 10 mM sodium acetate (pH 4.2) to 2.0 µM and immobilized on a CM5 sensor chip using random amine coupling kit (GE Healthcare) to a density of 3077 response units (RU) and the Sensorgrams were recorded at 25°C in the same buffer used for the ITC (including 150 mM NaCl and 0.005% P20 surfactant) and analyzed as described (24). Experiments were performed in independent duplicates in the range 3.42 nM–2500 nM for LNB, GNB, Lewis A triose and Blood group H antigen triose. Binding was also tested using 0.1 µM–100 mM Lac, Gal, 3FL, 3SL, DFL LNT, and di-*N*-Acetyl-lactosamine.

### Crystallization and structure determination of Blon_0883

Crystals were only obtained in the presence of either 1 mM GNB, 1 mM LNB or 8 mM Le^b^ by vapor diffusion in hanging drops and grew for one week at room temperature at a 1:1 ratio Blon_0883 (26 mg mL^-1^ in 10 mM MES pH 6.5 and 150 mM NaCl) and reservoir solution (0.2 M NaCl, 0.1 M Tris pH 7.0, 30% PEG3000 + TCEP. For the co-crystallisation with Le^b^, 1 μL protein: 1.5 μL reservoir was used. The crystals were flash frozen in liquid nitrogen without cryo-protectant. Diffraction data were collected to a maximum resolution of 2.1 Å and 1.4 Å for Blon_0883 complexed with LNB and GNB, respectively, at the Swiss Light Source, Paul Scherrer Inst, Villigen, Switzerland. Diffraction data were collected to a maximum resolution of 2.7 Å for Blon_0883 co-crystallised with the H-1 trisaccharide, respectively.

All datasets were processed with XDS (37). The structure was solved in the Orthorhombic space group *P2*_1_*2*_1_*2*_1_ using native-SAD phasing at 2.075 Å, data was collected as described by Weinert *et. al.* (38). The program Phenix.AutoSol (38–40) was used for solving the phases. An initial partial model was obtained with Phenix.AutoBuild (41). Further corrections and model building were performed using the program Coot (42) resulting in a complete model, which was used in molecular replacement to solve the structure of Blon_0883 in complex with LNB. The models were refined using phenix.refine (43) randomly setting aside 5% of the reflections. Molecular replacement with the protein part of Blon_0883 was used to solve the GNB-complexed and H1-complexed structures. Ligand molecules were built manually using Coot after the protein atoms were built and water molecules were added. The overall quality of all models was checked using MolProbity (44). Data collection and refinement statistics are shown in Supplementary Table S2. The two complexes are very similar in conformation and superposition of the individual models results in pairwise overall RMSD of 0.24 Å between aligned C_α_ atoms. The PyMOL Molecular Graphics System, Version 2.5.5 (Schrödinger, LLC, NY) was used to explore the models and for molecular graphics.

### MD Simulations

The binding of H1 and Le^a^ fucosylated trisaccharides to Blon_0883 was studied at atomic level using molecular dynamics (MD) simulations. The complexes were investigated over a 200 ns time scale in aqueous solution. Following the MD simulations, the interactions between the trisaccharides and the protein were analyzed. The docking procedure and MD simulation setup are briefly described (Detailed description is provided in the supporting information)

#### Docking and preparation of the systems

The structures of the H1 and Le^a^ trisaccharides were generated using GLYCAM (https://glycam.org/), ensuring correct saccharide nomenclature and alignment with the force field used for the MD simulation.

The crystal structure of the H1 complex was used to prepare the ligand-protein system for molecular dynamics (MD) simulations using the Protein Preparation Wizard in the Maestro Schrödinger Suite(45, 46). The ligands were prepared using the Ligand Preparation tool (47) in the Schrödinger Suite 2022. These ligands were then docked to Blon_0883 using Glide(48, 49). The docking procedure was validated by re-docking the H1 ligand statically to Blon_0883. For the Le^a^ ligand, a flexible docking procedure was employed, exploring all possible ligand conformations within the binding pocket. Since there is no crystal structure for the Le^a^-Blon_0883 complex, two binding modes were selected. The first mode mimics the binding of the H1 ligand. In the second mode, the dihedral angle between the Gal and GlcNAc units was rotated, flipping the GlcNAc ring 180 degrees. This repositioned the Fuc unit in the same pocket, where the fucose unit of the H1 antigen trisaccharide in complex with the protein. near the binding pocket’s opening. Each ligand-protein complex was processed using H++ (http://biophysics.cs.vt.edu/H++, version 4.0)(50–52) to ensure correct protonation of titratable residues at pH 7.0 and 0.15 M NaCl.

#### Molecular dynamic simulations

The MD simulations were carried out in Amber19.17(53). Topology and coordinate files were generated with AMBERTOOLS, and glycan/protein force fields (GLYCAM_06j-1, ff99SB) were assigned using LEaP(54–56). The systems were solvated in a truncated octahedral box with TIP3P water [14], neutralized, and ionized to match a 0.15 M salt concentration. MD simulations were performed in Amber19 applying the SHAKE constraint (57) that enables a 0.002 ps timestep. The number of steps during minimization varied to generated different initial conditions, followed by heating to 295.15K in the *NVT* (constant number of atoms, *N*, constant volume, *V*, and constant temperature, *T*) ensemble, with restraints on protein backbone and key hydrogen bonds. After equilibration in the *NpT* (constant number of atoms, *N*, constant pressure, *p*, and constant temperature, *T*) ensemble (58, 59), 200 ns constant pH MD simulations were performed in the *NpT* ensemble. During minimization and MD simulations, a 12 Å cut-off was used for non-bonded interactions, and the particle mesh Ewald method (60) for electrostatics. Coordinates were saved every 10 ps, and MD trajectories were analyzed to assess conformational changes and key interactions between ligands and Blon_0883. Ligand binding energies were determined using the Molecular Mechanics Generalized Born Surface Area approach (61) in cpptraj (53, 61).

### Genomic conservation and phylogenetic analysis of Blon_0883 homologs

Sequences were fetched from NCBI blastp (https://.ncbi.nlm.nih.gov/, 19^th^, September 2024), using the biochemically and structurally characterized Bifidobacterium SBP sequences lacking signal peptides as queries and limiting the search to sequences *Bifidobacterium* (taxid: 1678) with a lower amino sequence identity threshold of 30 % sequence identity to the query. Redundancy was reduced using CD-HIT (accesses through a local server installed from https://github.com/weizhongli/cdhit-web-server) with a 95 % sequence identity cutoff. Alignment was performed using MAFFT V. 7, and phylogenetic tree constructed using the Neighbor-joining algorithm with bootstraps performed (1000 iterations) on the MAFFT (62) server.

### Bacteria and culture conditions

*B. infantis* JCM 1222^T^ was obtained from Japan Collection of Microorganisms (RIKEN BioResource Center, Japan). *B. infantis* was anaerobically grown at 37°C in Gifu anaerobic medium (GAM; Nissui Pharmaceutical, Tokyo, Japan) or in basal medium (pH 6.7) composed of 0.2 % yeast extract, 1.0 % peptone, 0.5 % sodium acetate, 0.02 % MgSO_4_, 0.2 % K_2_HPO_4_, 0.18 % cysteine hydrochloride, 0.11 % NaCO_3_ (all w/v), 0.1 % (v/v) Tween80, as well as 0.5 % (w/v) of Gal, LNB, or GNB as a sole carbon source. Where appropriate, chloramphenicol (Cm; 2.5 μg mL^-1^) and spectinomycin (Sp; 10 μg mL^-1^) were used. The anaerobic culture was carried out in InvivO_2_ 400 workstation (Ruskinn Technology, Bridgend, UK; 10 % CO_2_, 10 % H_2_, and 80 % N_2_). Growth was monitored by measuring optical density at 600 nm (*OD*_600_).

### Targeted disruption of *Bi*GFL-BP gene (Blon_0883) in *B. infantis*

The *Bi*GFL-BP gene (Blon_0883) of *B. infantis* JCM 1222^T^ was inactivated by single-crossover-mediated insertional mutation using the method described previously(21). *Escherichia coli* DH5α was used as a host for plasmid construction. The homologous regions (internal region of *Bi*GFL-BP gene) were amplified by PCR using *B. infantis* genome as a template. The primers used are 5’-<u>CCAGCTCAAGGGATC</u>TGACCACATCCAAGAAGATC-3’ and 5’-<u>CGGTACCCGGGGATC</u>CACGTCCTTCTTGTAGGTCA-3’, with the 15-bp extension required for In-Fusion cloning underlined (Clontech Laboratories, Mountain view, CA, USA). The resulting PCR fragments were ligated within the BamH-digested 2.0 kb fragment of pBS423 (63), which was provided from RIKEN BRC through the National BioResource Project of the MEXT/ANED, Japan. After sequence confirmation, the plasmid was introduced by electroporation into the wild-type strain for genomic integration. Inactivation of the *Bi*GFL-BP gene (Blon_0883) was also conducted with the ΔBlon_2175–2177 basis (an orthologue of the *B. longum* GNB/LNB (GL-BP) transporter (Sakanaka *et al*., *in revision*)) (21) to generate a double KO strain. All of the strains including wild-type were made Cm-resistant by transformation with pBSF109 (21) so that the genotypes are the same except the two SBP genes.

### Quantification of LNB

LNB concentration in the spent media was analysed using high-performance liquid chromatography with a charged aerosol detector (HPLC-CAD; Thermo Fischer Scientific, Waltham, MA, USA) as described previously(64). This analysis was performed at 40°C using HILICpak VG-50 4E column (4.6 × 250 mm, Showa Denko K.K., Tokyo, Japan). Elution was performed with 73% acetonitrile for 30 min at a flow rate of 1.0 mL min^-1^. The standard curve was created using the known concentrations of LNB.

### Thin-layer chromatography for sugar consumption analysis

WT and mutant *B. infantis* cells cultivated overnight in liquid GAM were harvested by centrifugation and washed twice with the sugar-free basal medium. Thereafter, cells were suspended in the basal medium containing 5 mM Le^a^ or H1 trisaccharides as the sole carbon source to an *OD*_600_ of 2.0. The samples were taken from the cultures at the indicated time points, and the supernatants were collected by centrifugation. Sugars were analyzed by thin-layer chromatography (Silica gel 60, Sigma-Aldrich) using a solvent system of 1-butanol, acetic acid, and water (2:1:1 by volume) and visualized as described previously(65). A similar procedure was used to analyze the depletion of the Le^b^ tetrasaccharide, with the difference that *B. infantis* cells were grown on YCFA medium supplemented with 0.5 % (w/v) of a home preparation of complex HMOs from mothers’ milk in preculture and then 20 mL culture in the same medium and complex HMO concentration for 8 h. Then the cells were harvested and washed them 3 times in phosphate buffered saline, pH=6.8 and resuspended to an *OD*_600_ = 6 in the same buffer and a final substrate concentration of 2 mM Le^b^ tetrasaccharide. Aliquots of either 1 or 3 µL were collected at 0, 1h, 2h, 4h, 6h, and overnight and spotted on TLC plates together with Le^b^ standard. Sugars were separated using a mobile phase of 1-butanol, ethanol, and water (5:3:2 by volume) and visualized by spraying with 2 % (w/v) 5-methylresorcinol, 80 % (v/v) EtOH, and 10 % (v/v) H_2_SO4 and tarring at 300 °C.

## SUPPLEMENTAL MATERIAL

Supplemental material is available online xxxxx

Figures S1 to S7, Supplementary Tables S1-S5, Molecular Dynamic simulation details.

ASM does not own the copyrights to Supplemental Material that may be linked to, or accessed through, an article. The authors have granted ASM a non-exclusive, world-wide license to publish the Supplemental Material files. Please contact the corresponding author directly for reuse.

## DATA AVAILABILITY

The coordinate structural files have been deposited in the Protein Data Bank (PDB) under the accessions 9H0N, 9H0O and 9H0P.

## ACKNOWLEDGMENTS

We thank Motomitsu Kitaoka (Niigata University) for the kind gift of LNB and GNB, Satoru Fukiya and Atsushi Yokota (Hokkaido University) for helpful suggestions for *Bifidobacterium* gene disruption, Yuta Sugiyama (Ishikawa Prefectural University) for the support of LNB concentration analysis (HPLC-CAD analysis). We would like also to thank Associate Professor Albert Guskov for help with the Data collection.

## AUTHOR CONTRIBUTION

ME performed the cloning, binding, initial crystallization of Blon_0883 with LNB and GNB. HS performed the crystallization and determined the structure of Blon_0883 in complex with the H1 trisaccharide with PM. MJ completed the refinement of the structures, performed structural comparisons, and deposited the coordinate files. MJ also performed the phylogenetic analyses and made the structural figures and made the crystallography and structural comparison Tables. MS and SM constructed *B. infantis* mutant strains and conducted growth experiments and sugar concentration analysis using HPLC. MS and AN were responsible for the sugar consumption analysis using TLC. MJP performed the growth and Le^b^ uptake experiment. SK performed the initial parametrization and GHJP and JFH performed additional MD analyses and made the figures related to the MD work. The first manuscript draft was written by ME and MAH. TK and MS contributed to the early drafts and gave feedback on the later versions of the manuscript. All authors contributed to the MS and approved the final version.

